# Targeting Stem-loop 1 of the SARS-CoV-2 5’UTR to suppress viral translation and Nsp1 evasion

**DOI:** 10.1101/2021.09.09.459641

**Authors:** Setu M. Vora, Pietro Fontana, Valerie Leger, Ying Zhang, Tian-Min Fu, Judy Lieberman, Lee Gehrke, Ming Shi, Longfei Wang, Hao Wu

**Author notes:** Correspondence to Hao Wu; Ming Shi; or Longfei Wang. S.M. Vora, P. Fontana, M. Shi and L. Wang contributed equally to this paper.

## Abstract

SARS-CoV-2 is a highly pathogenic virus that evades anti-viral immunity by interfering with host protein synthesis, mRNA stability, and protein trafficking. The SARS-CoV-2 nonstructural protein 1 (Nsp1) uses its C-terminal domain to block the mRNA entry channel of the 40S ribosome to inhibit host protein synthesis. However, how SARS-CoV-2 circumvents Nsp1-mediated suppression for viral protein synthesis and if the mechanism can be targeted therapeutically remain unclear. Here we show that N- and C-terminal domains of Nsp1 coordinate to drive a tuned ratio of viral to host translation, likely to maintain a certain level of host fitness while maximizing replication. We reveal that the SL1 region of the SARS-CoV-2 5’ UTR is necessary and sufficient to evade Nsp1-mediated translational suppression. Targeting SL1 with locked nucleic acid antisense oligonucleotides (ASOs) inhibits viral translation and makes SARS-CoV-2 5’ UTR vulnerable to Nsp1 suppression, hindering viral replication in vitro at a nanomolar concentration. Thus, SL1 allows Nsp1 to switch infected cells from host to SARS-CoV-2 translation, presenting a therapeutic target against COVID-19 that is conserved among immune-evasive variants. This unique strategy of unleashing a virus’ own virulence mechanism against itself could force a critical trade off between drug resistance and pathogenicity.

---

Severe Acute Respiratory Syndrome Coronavirus 2 (SARS-CoV-2), the causative agent of the infectious disease COVID-19, is a highly contagious and deadly virus with fast person-to-person transmission and potent pathogenicity (Chan et al., 2020a; Chan et al., 2020b). It is an enveloped, single-stranded betacoronavirus that contains a positive-sense RNA genome of about 29.9 kb (Lu et al., 2020; Wu et al., 2020; Zhu et al., 2020). The SARS-CoV-2 genome codes for two large overlapping open reading frames (ORF1a and ORF1b) and a variety of structural and nonstructural accessory proteins (Zhou et al., 2020). Upon infection, the polyproteins ORF1a and ORF1b are synthesized by host machinery and proteolytically cleaved into 16 mature non-structural proteins, namely Nsp1 to Nsp16 (Chan et al., 2020a; Chan et al., 2020b; Hartenian et al., 2020; Zhou et al., 2020).

Nsp1 is a critical virulence factor of coronaviruses and plays key roles in suppressing host gene expression, which facilitates viral replication and immune evasion, presumably by repurposing the host translational machinery for viral production and preventing the induction of type I interferons (Jimenez-Guardeno et al., 2015; Kamitani et al., 2009; Lei et al., 2020; Tanaka et al., 2012). It has been shown that Nsp1 effectively suppresses the translation of host mRNAs by directly binding to the 40S small ribosomal subunit (Narayanan et al., 2008; Tohya et al., 2009). Recent cryo-electron microscopy (cryo-EM) structures of SARS-CoV-2 Nsp1 indeed reveal the binding of its C-terminal domain (CT) to the mRNA entry channel of the 40S subunit (Harvey et al., 2021; Schubert et al., 2020) which contributes to blocking translation. These structural data are further supported by experiments demonstrating that Nsp1 binding to the 40S ribosome requires an open head conformation induced by core elongation initiation factors and that Nsp1 cannot bind to a 40S with an mRNA already occupying the entry channel (Lapointe et al., 2021).

Besides directly inhibiting mRNA translation, Nsp1 has also been shown to reduce the available pool of host cytosolic mRNAs by both promoting their degradation and inhibiting their nuclear export (Finkel et al., 2021; Simeoni et al., 2021; Zhang et al., 2021). Mutants of Nsp1 that disrupt ribosome binding also abolish mRNA degradation, suggesting that the degradation is likely downstream of Nsp1 translational block, and these two processes likely synergize to shut off host protein expression (Huang et al., 2011a; Narayanan et al., 2008).

Previous studies of SARS-CoVs have also implicated the 5’ untranslated region (5’ UTR) of coronavirus mRNAs in protecting the virus against Nsp1-mediated mRNA translation inhibition (Huang et al., 2011b; Kamitani et al., 2009; Thoms et al., 2020). However, how SARS-CoV-2 overcomes Nsp1-mediated translation suppression for its replication and whether this mechanism can be targeted for therapeutic intervention remain open questions. Here, we reveal that SARS-CoV-2 uses the short stem-loop 1 (SL1) in its 5’ UTR to escape Nsp1 suppression to effectively switch the translational machinery from synthesizing host proteins to making viral proteins, and that both the CT and the N-terminal domain (NT) are required for the transition from host to viral translation. The latter is supported by complementary experiments in a preprint released while the manuscript was under preparation {Mendez, 2021 #31}. We further show that SL1 can be targeted by locked nucleic acid (LNA) antisense oligonucleotides to prevent SARS-CoV-2 5’ UTR from evading its own translational suppression to potently inhibit viral replication.

## RESULTS

### SARS-CoV-2 5’ UTR mediates translation despite the presence of Nsp1

To investigate the function of SARS-CoV-2 Nsp1 in inhibiting mRNA translation, we co-transfected an mScarlet reporter construct with MBP-tagged Nsp1 or the MBP control in HeLa cells, and imaged mScarlet fluorescence and anti-MBP immunofluorescence (Fig. 1 A and B). The mScarlet reporter used an expression vector that contains the cytomegalovirus (CMV) promoter and 5’ UTR, and is commonly employed for mammalian cell expression (referred to as control reporter throughout the manuscript). The MBP and MBP-Nsp1 constructs also used the CMV promoter and 5’ UTR. Upon analysis of the control reporter, we found that mScarlet expression in MBP-Nsp1-transfected cells was reduced by over 7.1-fold (p<0.001) compared to cells co-transfected with MBP-alone (Fig, 1 B and C). These data are consistent with previous reports indicating potent translational suppression by Nsp1 (Schubert et al., 2020; Thoms et al., 2020).

**Figure 1.**
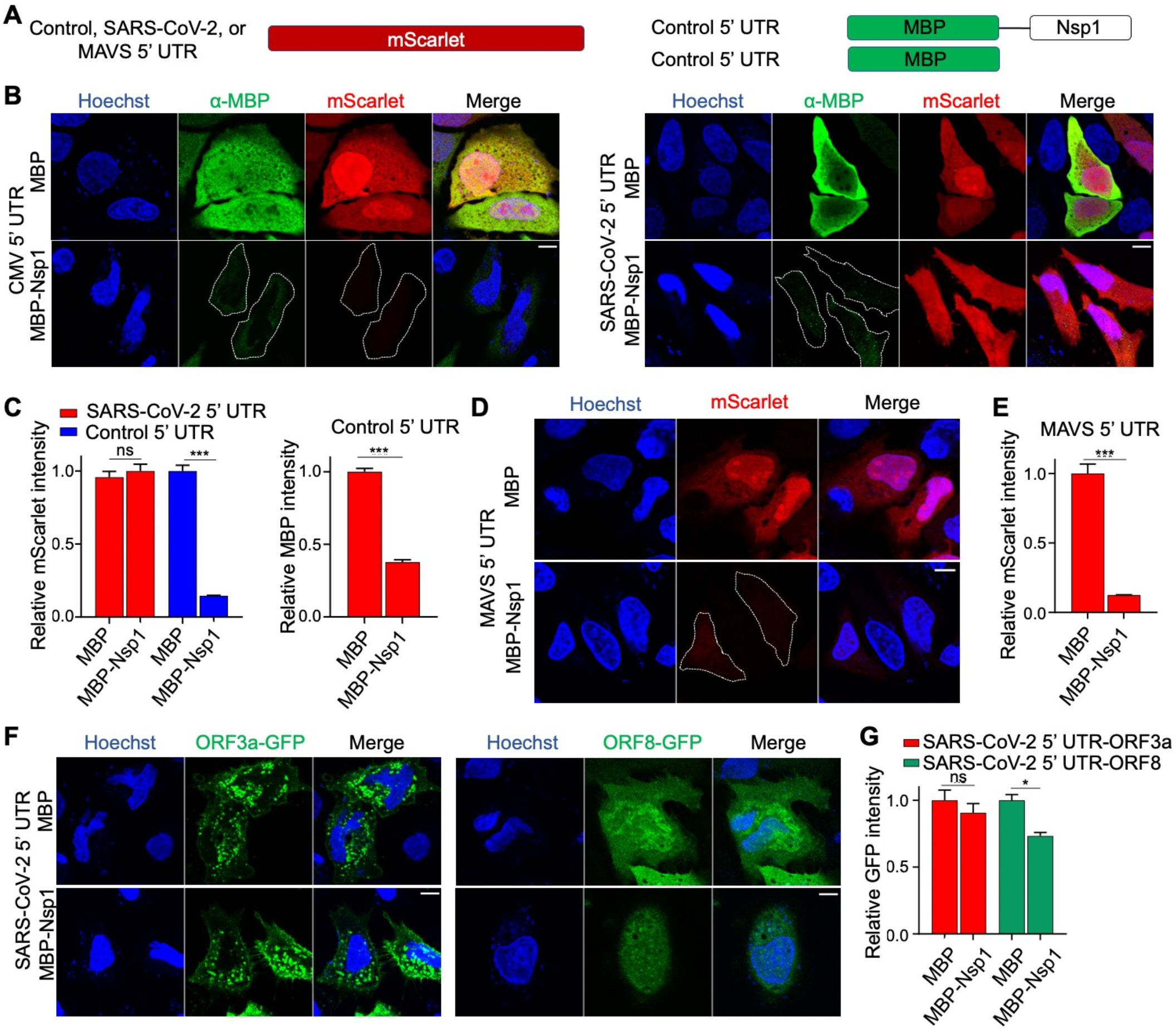
SARS-CoV-2 5’ UTR bypasses Nsp1-mediated inhibition of translation. (A) Schematic of translational reporters. 5’ UTR sequences from Control, MAVS, or SARS-CoV-2 were placed upstream of the mScarlet reporter (left panel). MBP or MBP-Nsp1 (right panel) were both downstream of Control 5’ UTR and were co-transfected along with each reporter plasmid. A CMV promoter was used to drive expression in all constructs. (B) Representative images of HeLa cells co-transfected with Control 5’ UTR reporter or SARS-CoV-2 5’ UTR reporter and either MBP alone or MBP-Nsp1 and visualized for DNA by Hoechst (blue), MBP by indirect immunofluorescence (green), and mScarlet by *in situ* fluorescence (red). Successfully transfected cells difficult to visualize due to low intensity are outlined here and in other figures. (C) Quantification of relative mScarlet intensity of data corresponding to (B). (D) Representative images of HeLa cells transfected with MAVs 5’ UTR reporter (red). (E) Quantification of relative mScarlet intensity in (D). (F) HeLa cells transfected with either ORF3a-GFP (left panel) or ORF8-GFP(right panel) downstream of SARS-CoV-2 5’ UTR. (G) Quantification of relative GFP intensity in (F). Error bars correspond to standard error of the mean except where otherwise noted. Scale bars are shown in each bottom right image and correspond to 10 microns.

On the other hand, when the 5’ UTR in the control mScarlet reporter was replaced by the SARS-CoV-2 5’ UTR (referred to as CoV-2 reporter throughout the manuscript), no significant difference in mScarlet expression was observed upon co-expression with MBP-Nsp-1 or MBP, indicating robust evasion of Nsp1-mediated translational suppression (Fig. 1 A-C). By contrast, in this same experiment, CMV 5’ UTR-controlled MBP-Nsp1 showed significantly lower expression than MBP alone (p<0.001) (Fig. 1 B and C). Thus, expression of Nsp1 from this construct was likely self-limiting, yet still sufficient to inhibit translation of mRNAs with non-SARS-CoV-2 5’ UTR but not the reporter with SARS-CoV-2 5’ UTR. Since the CMV 5’ UTR is not representative of those in human mRNAs, we also generated a reporter containing the 5’ UTR from human MAVS, an essential signalling effector responsible for certain virus-induced production of type I and III IFNs including SARS-CoV-2 (Yin et al., 2021). This mScarlet reporter was also potently suppressed by MBP-Nsp1, which decreased its expression 8.1-fold relative to MBP alone (p<0.001) (Fig. 1 A, D and E), suggesting that Nsp1-medaited translational suppression of host mRNAs can contribute to disabling critical mediators of the anti-viral IFN response.

To model subgenomic RNAs generated during discontinuous SARS-CoV-2 gene transcription, we tested constructs with SARS-CoV-2 5’ UTR upstream of either GFP-fused ORF3a or ORF8 (ORF3a-GFP or ORF8-GFP). ORF3a-GFP expression was not significantly decreased by Nsp1 co-expression, and ORF8-GFP showed only a 1.4-fold decrease (Fig. 1 F and G), which was modest relative to the 7.1-fold and 8.1-fold decrease seen with CMV 5’ UTR and MAVS 5’ UTR, respectively (Fig. 1 C and E). Together these data support that SARS-CoV-2 Nsp1 potently inhibits host protein translation and that SARS-CoV-2 5’ UTR allows evasion of Nsp1-mediated suppression.

### The SL1 of the 5’ UTR is necessary and sufficient for evasion of Nsp1-mediated translation suppression

The 5’ UTR of coronaviruses comprises a number of stem-loop structures (Fig. S1), among which SL1 has been shown to play critical roles in driving viral replication (Li et al., 2008; Narayanan et al., 2008; Schubert et al., 2020; Thoms et al., 2020; Tohya et al., 2009). For SARS-CoV-2, this conclusion should also make sense as the leader sequence driving all subgenomic RNAs is comprised of just SL1-SL3 instead of the entire 5’ UTR (Alexandersen et al., 2020; Kim et al., 2020), highlighting the potential importance of SL1 in both viral replication and possibly in promoting evasion from translation suppression by Nsp1. To test the latter function of the SL1 sequence of SARS-CoV-2, we generated ΔSL1 and SL1-alone 5’ UTR mScarlet reporters (Fig. 2 A). Compared with mScarlet translation in control cells co-transfected with MBP, the translation of SARS-CoV-2 ΔSL1 5’ UTR mScarlet reporter in MBP-Nsp1 transfected cells was 6.3-fold reduced, similar to our control reporter and MAVS reporter above (*p* < 0.001) (Fig. 2 B and C), a reduction similar to the control and MAVS 5’ UTR reporters (Fig. 1 C and E). These data suggest that SL1 is completely required for evasion of Nsp1-mediated translation suppression. Interestingly, the reporter bearing only the SL1 sequence in its 5’ UTR was not significantly reduced upon co-expression with Nsp1, indicating that the SL1 sequence is both necessary and sufficient for evasion of Nsp1-mediated translation suppression in our experimental system (Fig. 2 B and C).

**Figure 2.**
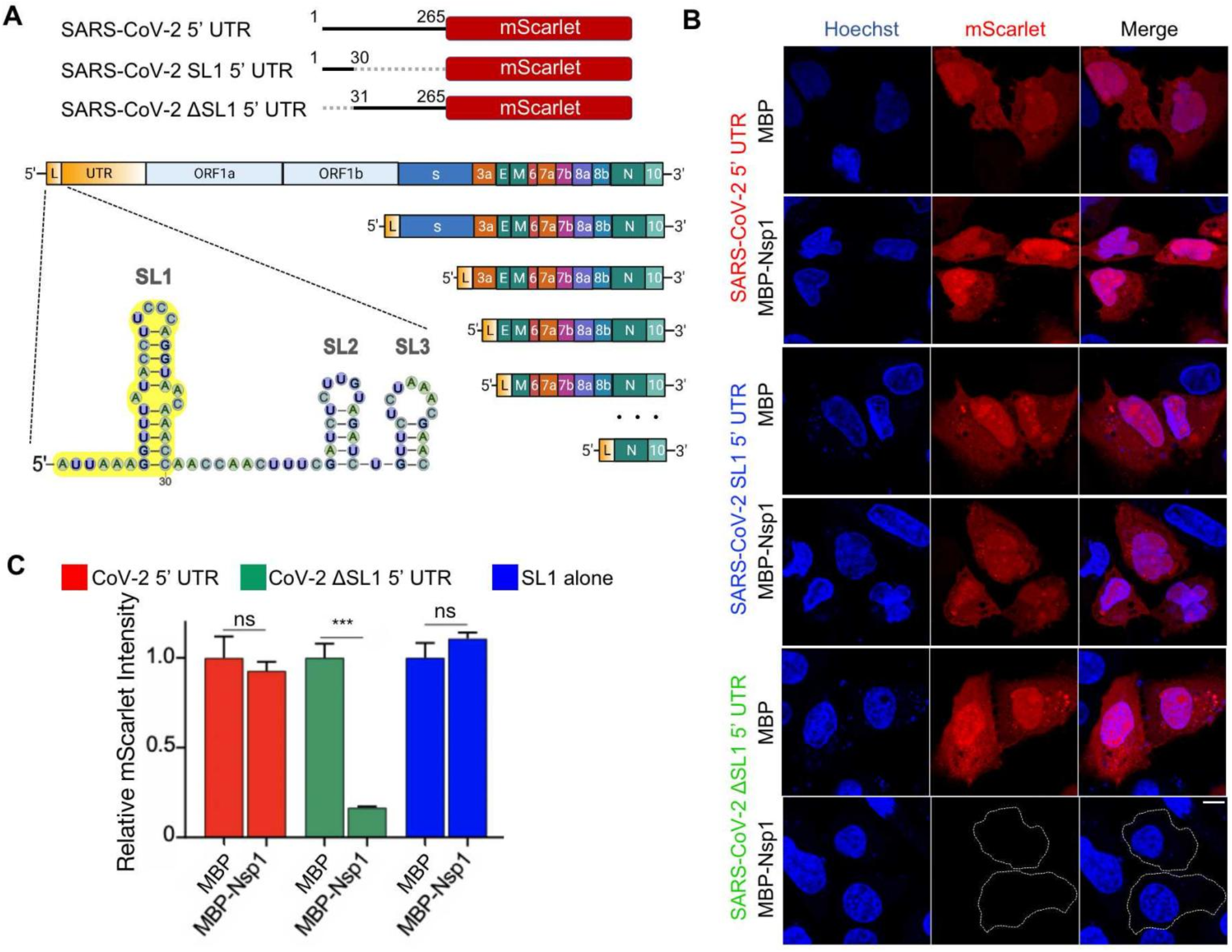
The SL1 stem-loop of the 5’ UTR is necessary and sufficient for evasion of Nsp1-mediated translation suppression. (A) Schematic representation of 5’ UTR, SL1 5’ UTR and ΔSL1 5’ UTR placed upstream of mScarlet (top panel), and of SARS-CoV-2 leader sequence containing SL1 (yellow) along with its incorporation into subgenomic RNAs (bottom panel). (B) Representative images of HeLa cells co-transfected with SARS-CoV-2 5’ UTR reporter and either MBP alone or MBP-Nsp1, and visualized for DNA by Hoechst (blue) and mScarlet by *in situ* fluorescence (red). (C) Quantification of relative mScarlet intensity of data corresponding to (B).

### Nsp1-CT and Nsp1-NT are both required for optimal host suppression and SL1-driven bypass

Nsp1 is a 187 amino acid (aa) protein with a 128 aa N-terminal domain and a 33 aa C-terminal domain separated by a short linker region (Fig. 3 A). Previous structures of the Nsp1− ribosome complex suggested that Nsp1-CT blocks mRNA entry to the ribosome and should be sufficient to inhibit protein translation (Schubert et al., 2020; Thoms et al., 2020). In order to probe the relative functions of these domains in suppressing host translation and allowing bypass by SARS-CoV-2 5’ UTR, we co-transfected HeLa cells with SARS-CoV-2 or control reporters along with Nsp1 FL, NT, CT, or NT+CT constructs (Fig. 3 B and C). None of these treatments significantly compromised CoV-2 5’ UTR reporter activity relative to FL Nsp1; however, NT, CT, and NT+CT less efficiently inhibited the control reporter mScarlet fluorescence intensity relative to FL Nsp1, with NT being the least effective (Fig. 3 B and C, Fig. S2). We repeated this experiment in HEK293T cells and found that the CoV-2 5’ UTR reporter activity was significantly higher with co-expression of NT and significantly lower with CT in comparison with FL Nsp1, suggesting impaired evasion of CT-imposed translational block (Fig. 3 D and E, Fig. S2). While FL Nsp1 most effectively suppressed the control reporter, the trend of suppression of the control reporter by NT, CT, or NT+CT mirrored that of the CoV-2 5’ UTR reporter (Fig. 3 E).

**Figure 3.**
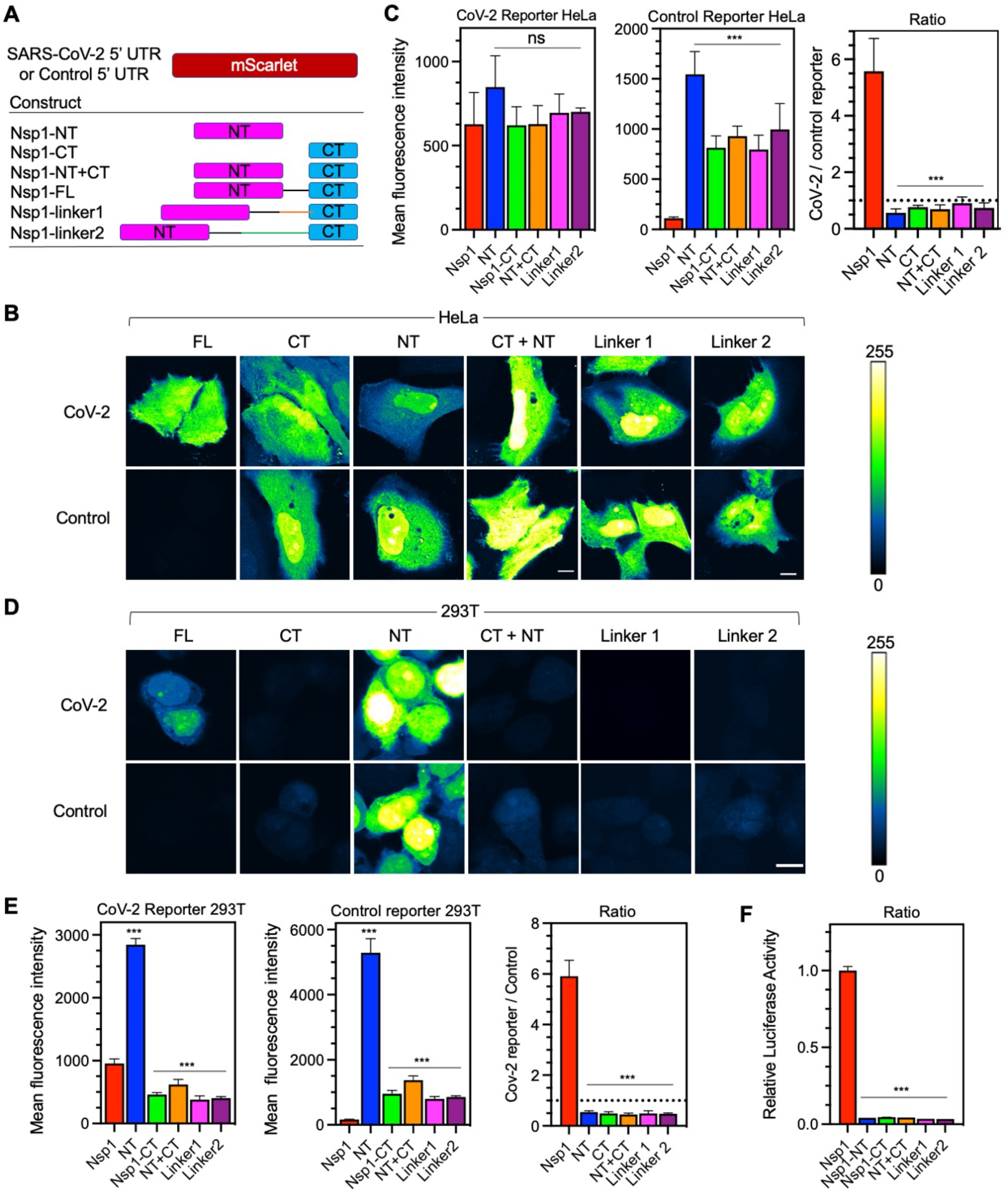
Nsp1 N- and C-terminal domains cooperate to drive viral translation selectivity. (A) Schematic of co-expression system with CoV-2 or control reporter along with various fragments of Nsp1 (FL, NT, CT, NT+CT) or extended linker mutants (linker1, linker2). (B) mScarlet fluorescence intensity in HeLa cells co-transfected with either the CoV-2 (top row) or control reporter (bottom row) along with various mutants of Nsp1. Intensity values are false coloured according to a scale (right panel). (C) Quantification of fluorescent intensity in (B) of CoV-2 (left) or control reporters (middle) and the ratio of CoV-2/Control (right) with different Nsp1 mutants. (Dashed line marks ratio of 1). (D) mScarlet fluorescence intensity in 293T cells as in (B). (E) Quantification of (D). (F) Relative luciferase activity in 293T cells assay co-transfected with CoV-2 firefly luciferase and control renilla luciferase reporters along with various Nsp1 mutants. The ratio of Firefly / Renilla luciferase was normalized and plotted in 3 replicates. Scale bars for panels (B) and (D) both correspond to 10 microns. Error bars represent standard deviation.

These apparently different observations from HeLa versus 293T cell lines were intriguing, but when we looked at the ratio of the CoV-2 5’ UTR reporter to the control reporter fluorescence, we found a strikingly similar trend. FL Nsp1 co-expression led to a CoV-2/control fluorescence ratio of 5.6 ± 1.2 and 5.9 ± 0.62 for HeLa and 293T cells, respectively (Fig. 3 C and E). In both cell lines, co-expression of either CT or NT led to a ∼10-fold reduction in CoV-2/control fluorescence ratio, and co-expression of NT+CT from separate constructs reduced CoV-2/control ratio to a similar extent (Fig. 3 C and E). To further validate these observations, we utilized a luciferase reporter in 293T cells -which also relies on ratiometric normalization to a control reporter - and found a similar reduction in CoV-2 reporter translation selectivity by the different Nsp1 constructs (Fig. 3 F). Collectively, these data suggest that NT and CT are both important for host translational suppression, which is consistent with a recent preprint (Mendez et al., 2021), and that the NT is required for viral evasion of Nsp1-mediated translational suppression. In addition, Nsp1 may be tuned to control the ratio of viral/host translation rather than simply promoting high viral translation, perhaps to maintain some level of host fitness to allow viral replication.

### Correct association and spacing of NT+CT via the Nsp1 linker is required for function

The fact that NT+CT expression from different constructs compromised CoV-2/control ratio suggested a role for covalent association between the two domains via a linker. To test whether the length of the linker between NT and CT in Nsp1 has any functional effect, we inserted an additional 20 residues (linker1) or 40 residues (linker2) at the Nsp1 linker region (Fig. 3 A). Remarkably, the linker extensions dramatically reduced the ratio between CoV-2 and control reporter expression (*p* < 0.001) in both HeLa and 293T cell lines (Fig. 3 B-E). These data were also validated in the SARS-CoV-2 5’ UTR luciferase assay (Fig. 3 F), suggesting that the NT and CT must somehow cooperate in a spatially specific manner to allow optimal suppression of host translation, and to permit the evasion of suppression on viral translation. Consistent with another report, we also observed lack of nuclear localization when visualizing Nsp1 FL, CT, linker1 and linker2 (Zhang et al., 2021). NT alone, however, showed both nuclear and cytosolic localization, suggesting a role for CT and/or linker regions for sequestering Nsp1 in the cytosol (Fig. S2).

Various naturally occurring mutations have been described in Nsp1 throughout the pandemic including a 3 amino acid deletion in the Nsp1 linker region (Nsp1^ΔKSF^) detected in North America and Europe (Benedetti et al., 2020; Lin et al., 2021). Given the importance of the Nsp1 linker length in regulating viral to host translation selectivity, we tested the function of this variant using our reporter assay. Although Nsp1^ΔKSF^ induced a small significant decrease in CoV-2 reporter activity, it more than doubled control reporter activity compared to WT Nsp1, and significantly reduced CoV-2/control translation ratio (p<0.001) (Fig. S3). Thus, while lengthening the linker alters regulation of both viral and host translation, the shortened linker in this variant mainly compromised suppression of host translation. Together, these results suggest that the Nsp1 linker length is optimized to coordinate host translational suppression and bypass by SARS-CoV-2 5’ UTR, and that the Nsp1^ΔKSF^ mutant could be less virulent.

It has been recently determined that Nsp1 promotes degradation of host mRNAs whose translation is suppressed (Finkel et al., 2021), which depends on R124, a key residue in the NT that is conserved in SARS-CoV (Kamitani et al., 2009; Thoms et al., 2020). To test whether this residue affects host translational shutdown, we co-expressed our control reporter with Nsp1^R124A^, which increased reporter activity by 5-fold relative to control (p<0.001). Interestingly, this mutation also led to a 1.5-fold decrease in CoV-2 reporter activity (p<0.001) and significantly reduced the CoV-2/Control ratio (p<0.001; Fig. S3). These results suggest that mRNA degradation could indeed contribute to host shutdown and is consistent with the idea that it reduces the pool of available host mRNAs able to compete with SARS-CoV-2 RNA for ribosome association. It was reported by two other studies that R124A does not interfere with translational suppression of host mRNAs, but these studies relied on in vitro translation in cell extracts that are not optimized to recapitulate Nsp1-directed host mRNA degradation (Lapointe et al., 2021; Mendez et al., 2021).

### SL1 antisense oligos (ASO) selectively target SARS-CoV-2 5’ UTR with nanomolar potency

To suppress viral translation, we attempted to disrupt SL1’s function using anti-sense oligonucleotides (ASO). Since SL1 is sufficient for both viral translation and evasion of Nsp1-mediated translation suppression, ASOs targeting SL1 could represent novel therapeutic opportunities to effectively inhibit viral translation. SL1 starts right after the 5’ cap, and its structure is dynamically regulated during viral replication (Ziv et al., 2020). The stem region of SL1 contains 10 Watson Crick base pairs with a bulge at the centre (Fig. 4 A) (Miao et al., 2021). We rationally designed different ASOs to hybridize with various regions of the SL1 and tested their activity against our CoV-2 5’ UTR reporter in the presence or absence of Nsp1.

**Figure 4.**
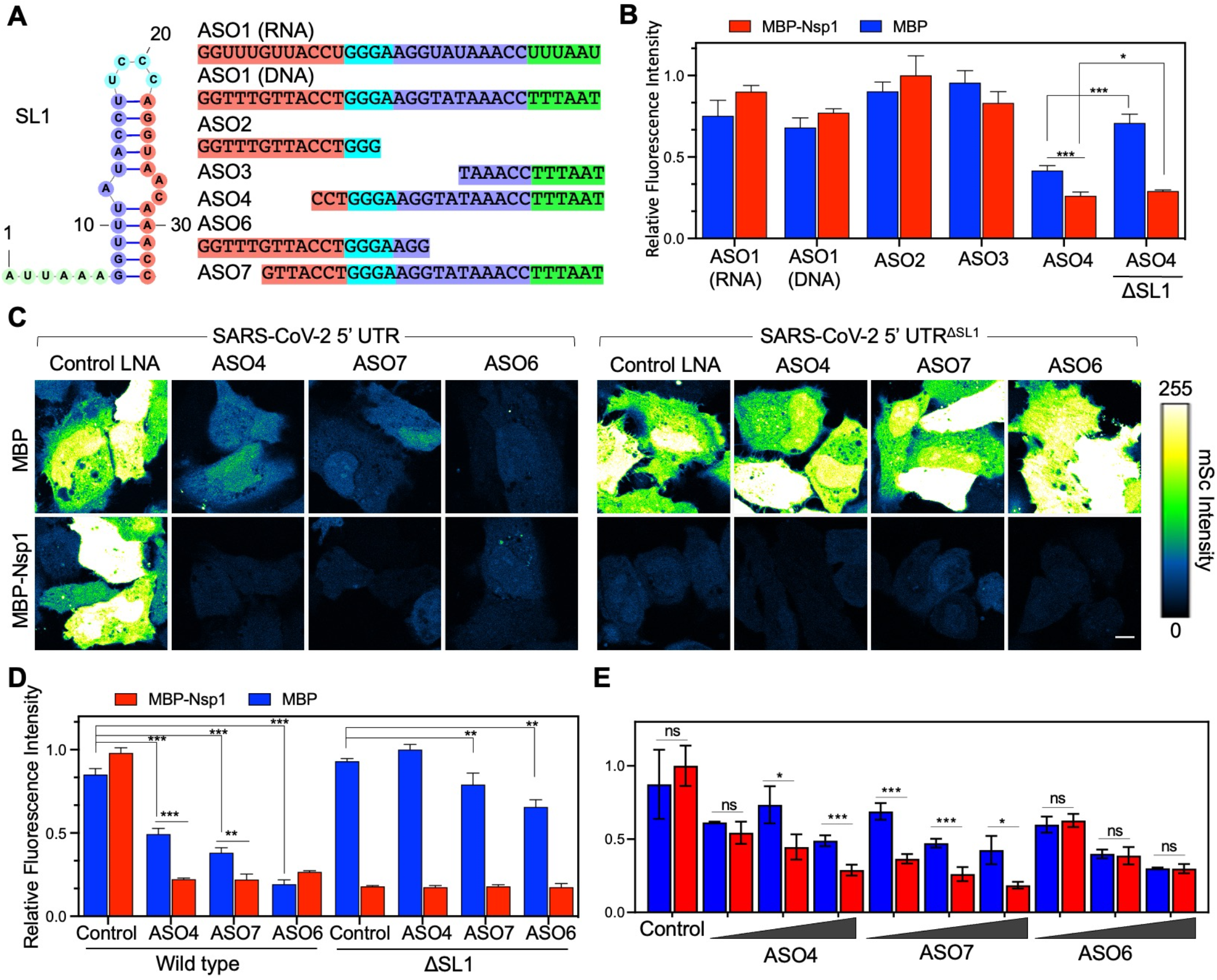
ASOs targeting SL1 renders the SARS-CoV-2 5’ UTR susceptible to Nsp1-mediated shutdown. (A) Schematic of SL1 region and various ASOs (which are all LNA miximers unless otherwise noted). Complementary sequences between ASOs and SL1 are matched by colour. (B) Initial screen of ASO activity. Each ASO was transfected at 50 nM with CoV-2 reporter along with either MBP alone or MBP-Nsp1. Bar on far right indicates co-transfection with a reporter lacking SL1 as a control. (C) Images of HeLa cells transfected with 50 nM ASO4, 7, 6, or a control ASO along with CoV-2 or ΔSL1 reporter and either MBP or MBP-Nsp1. Intensity values are false coloured according to a scale (right). (D) Quantification of (C). (E) Dose-response assay of each ASO. Cells were transfected with CoV-2 reporter and MBP or MBP-Nsp1 as above. ASOs were transfected at either 25, 50, or 100 nM.

In preliminary experiments, when transfected at 50 nM, neither DNA nor RNA anti-SL1 ASOs showed activity against CoV-2 5’ UTR in the presence or absence of Nsp1 (Fig. 4 B). We then further designed ASOs with locked nucleic acid (LNA) miximers targeting various regions of SL1 (Fig. 4 A). ASO2 and ASO3 LNAs (short, ≤ 15 bases) targeting the 5’ and 3’ regions of SL1, respectively, both failed to suppress SL1 activity (Fig. 4 B). ASO4, a 24 base LNA against the 3’ region of SL1, successfully suppressed reporter activity on its own when co-transfected at 50 nM with the CoV-2 reporter and MBP alone (Fig. 4 B). Interestingly, this suppression was further enhanced upon co-expression with Nsp1 (Fig. 4 B), indicating successful inhibition of SL1-mediated evasion of Nsp1 translational shutdown. In the presence of Nsp1, suppression of CoV-2 5’ UTR reporter activity by ASO4 was even significantly lower when compared to the ΔSL1 reporter, demonstrating that ASO4 induces a complete loss of function of the SL1 sequence (Fig. 4 B). We further designed 2 more LNA ASOs of similar lengths – ASO6 and ASO7 – against the 5’ and 3’ regions of SL1, respectively. Like ASO4, both ASO6 and ASO7 suppressed the SARS-CoV-2 5’ UTR reporter on their own and showed relatively little activity against the same reporter lacking the SL1 sequence (Fig. 4 C and D). Interestingly, only ASO4 and ASO7, but not ASO6, synergized with Nsp1 to further suppress SARS-CoV-2 5’ UTR activity (Fig. 4 C and D). This tendency was consistent when the ASOs were tested over various concentrations (25, 50, and 100 nM) (Fig. 4 E,). Together, these data suggest that ASO4 and ASO7 suppress viral translation in at least two ways: 1) by making the 5’ UTR less efficient in driving viral translation and 2) by interfering with the evasion of the 5’ UTR from Nsp-1 mediated suppression.

### SL1 antisense oligos effectively inhibit SARS-CoV-2 replication in Vero E6 cells

To test whether anti-SL1 ASOs with activity in reporter assays could also inhibit viral replication in Vero E6 cells, we first confirmed the function of these ASOs in this cell line by co-transfecting Vero E6 cells with SARS-CoV-2 5’ UTR reporter, Nsp1, and various ASOs. When co-transfected at 100 nM, ASO4 and ASO7 gave a 5-fold and 2-fold reduction, respectively, in reporter activity at 24 hours (Fig. 5 A). Importantly, this reduction in reporter activity persisted for at least 72 hours, indicating that they retained function and stability upon transient delivery to cells (Fig. 5 A). We then transfected Vero E6 cells with 100 nM of an ASO along with a control mScarlet reporter, followed by SARS-CoV-2 infection at either 0.1 or 0.5 multiplicity of infection (MOI). Cells were fixed 72 hours post-infection and stained for nucleocapsid (N) to mark infected cells (Fig. 5 B-E). Roughly 37% of Vero E6 cells were mScarlet^+^ when transfected with control ASO and mock infected, which was used to indicate transfection efficiency (Fig. 5 C). For either MOI 0.1 or 0.5, we observed a ∼4-fold reduction in N^+^ mSc^+^ cells with ASO4 and ASO6 and a ∼2-fold reduction with ASO7 when compared with control LNA (Fig. 5 C-E). Thus, our strategy for targeting SL1 with ASOs can successfully inhibit SARS-CoV-2 replication in vitro.

**Figure 5.**
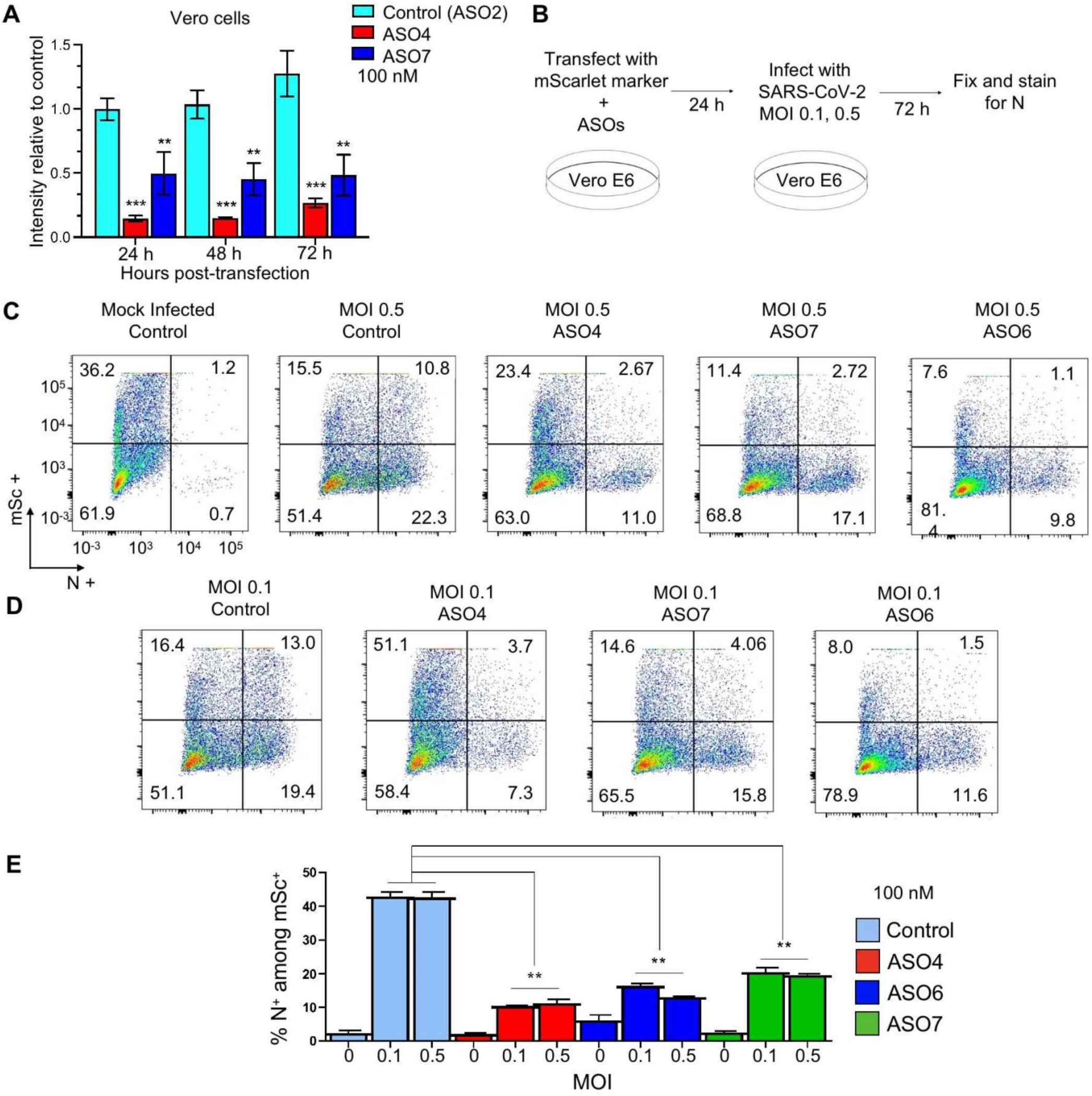
ASOs targeting SL1 produce stable knockdown after transient delivery and inhibit SARS-CoV-2 replication in vitro. (A) Various ASOs along with CoV-2 reporter and MBP-Nsp1 were transiently transfected into Vero E6 cells and reporter intensity was measured daily over the course of 72 hours, shown for each ASO (Control, ASO4, and ASO7) at each timepoint. Since expression from transfected plasmids naturally changes over time, each datapoint was normalized to intensity of a parallel control where no ASO was included. (B) Schematic of in vitro infection experiment. Vero E6 cells were co-transfected with an mScarlet (mSc) marker and various ASOs at 100 nM followed by infection with SARS-CoV-2 at MOI 0.1 or 0.5, and fixed and stained after 72 hours. (C) Nucleocapsid intensity plotted against mSc obtained by flow cytometry for each treatment (infected at MOI 0.5). Quadrants demarcate mSc+, N+ cells (top right quadrant), and the corresponding percentage of cells is listed in each corner. (D) Same as (C) except infection was at MOI 0.1. (E) Percent of successfully transfected cells (marked by mSc positivity) that were N+ by ASO treatment (colour coded according to legend on the right) at various MOIs. Error bars represent standard deviation.

## DISCUSSION

SARS-CoV-2 Nsp1 is a major virulence factor which suppresses host gene expression and immune defence (Jimenez-Guardeno et al., 2015; Kamitani et al., 2009; Tanaka et al., 2012). The recently published cryo-EM structures (Schubert et al., 2020; Thoms et al., 2020) showed that the helical hairpin at the CT of SARS-CoV-2 Nsp1 interacts with the 40S subunit of the ribosome to block mRNA entry. Here, we revealed that the NT also contributes to host translational suppression by coordinating with the CT, and that the NT and CT need to be covalently linked and correctly spaced to perform this function optimally. Insertion or deletion of linker residues at the NT to CT junction also compromises this function. We further found that SARS-CoV-2 5’ UTR evades Nsp1 suppression to allow viral translation, which also requires both NT and CT. This finding is consistent with our analysis on the Nsp1 NT-CT linker deletion mutant as well as with other naturally isolated SARS-CoV-2 NT deletion mutants which were associated with lower viral titres and less severe COVID-19 disease (Lin et al., 2021).

Our data show that the Nsp1 NT and CT coordinate to perform two functions: host translational suppression and bypass of this suppression by SARS-CoV-2 5’ UTR. The final outcome is that Nsp1 controls the viral to host translation ratio rather than simply promoting high viral translation, via both its NT and CT. Interestingly, while the effects of these two domains on absolute viral or host reporter activity differed by cell type, the ratio of viral to host translation was invariant. This tempered viral to host translation ratio imposed by Nsp1 could be optimal to maintain some level of host fitness to allow viral replication. It may also be tuned to produce the maximally allowable viral copy number that could still avoid triggering an interferon response. Thus, our studies show that coronaviruses like SARS-CoV-2 have evolved a clever strategy involving Nsp1 and 5’ UTR to control the host translation machinery to support viral replication to effectively counteract the host cellular immune system (Fig. 6 A).

**Figure 6.**
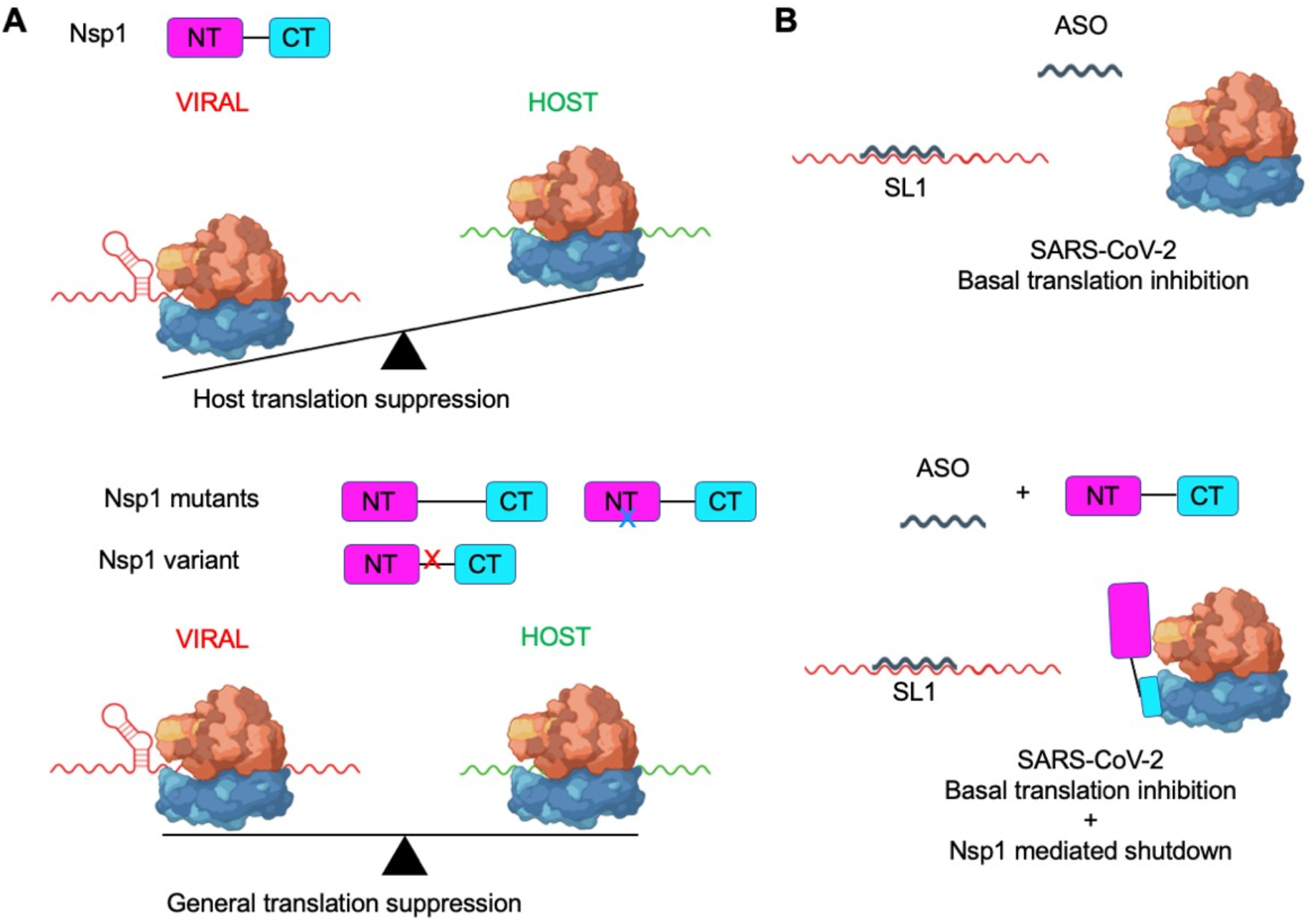
Model for Nsp1-driven viral translation selectivity and its disruption via ASO targeting of SL1. (A) Nsp1 shuts down host translation, mainly by blocking the mRNA entry channel of the 40S ribosome which ultimately results in host mRNA degradation. The SL1 region in SARS-CoV-2 5’UTR allows evasion of translational suppression, leading to selective viral translation (top panel). Removal of NT or CT domains - or disruption of their proximity by lengthening or shortening the linker region – all result in compromised host suppression and viral evasion, resulting in a reduced viral to host translation ratio (bottom panel). (B) Targeting SL1 via ASO can disrupt basal translation (top panel) and more importantly, makes SARS-CoV-2 5’UTR vulnerable to Nsp1-mediated translation suppression (bottom panel), effectively halting viral replication.

A remaining question is whether the major driver of Nsp1-mediated host shutoff is depletion of host mRNAs or direct inhibition of translation. A recent study using ribosomal profiling and RNA sequencing found similar translation efficiencies of viral and host mRNAs during SARS-CoV-2 infection of Calu3 cells (Finkel et al., 2021). One caveat is that these measurements were taken in late timepoints when viral RNAs had already started to enter the endomembrane system for packaging and would thus be shielded from the translation in the cytosol, skewing the number of elongating transcripts per copy(Finkel et al., 2021). A role for selective and direct translational block of host mRNAs is also supported by in vitro translation experiments (Mendez et al., 2021; TIDU et al., 2020). Nonetheless it is likely that both mRNA degradation and inhibition of translation are mechanistically and functionally linked and both are major contributors to SARS-CoV-2 virulence.

The exact molecular basis for how Nsp1 NT coordinates with SL1 of SARS-CoV-2 5’ UTR to bypass the translation inhibition is still not yet clear. A previous study suggested that both FL and ΔCT Nsp1 directly bind the SL1 region of the CoV-2 5’ UTR by gel shift (Vankadari et al., 2020). However, this study was performed under low salt conditions, and we could not detect such interactions under physiological salt concentrations (data not shown). Thus, the molecular details of NT coordination with SL1 remain elusive, and it is likely mediated by an indirect physical interaction. Future studies will be critical for teasing out the functional interactions between Nsp1 and host proteins.

Our data also demonstrates that a single cis-acting element in the SARS-CoV-2 5’UTR, the SL1 stem-loop, is both necessary and sufficient for evading Nsp1-mediated host shutdown and is thus a vulnerable therapeutic target for limiting SARS-CoV-2 replication (Fig. 6 B). We revealed that SL1-targeting ASOs, including ASO4, ASO6 and ASO7, potently suppress SARS-CoV-2 5’ UTR reporters in HeLa and Vero E6 cells, and SARS-CoV-2 replication in Vero E6 cells at nanomolar concentrations. Importantly, our ASOs were able to make SARS-CoV-2 vulnerable to its own mechanism of Nsp1-mediated host shutdown, a benchmark that could inform the development of future therapeutics targeting this critical mechanism. As we were preparing our manuscript, an independent preprint reported unbiased screening experiments which identified a promising ASO that is complementary to the 5’ UTR sequence, from the SL1-SL2 linker to the last 5 nucleotides of SL1, further underscoring its importance as a target (Zhu et al., 2021) (Fig. S4). However, it is unknown whether this ASO also sensitize SARS-CoV-2 to its own Nsp1-mediated translational suppression, like our ASO4 and ASO7.

Given that within the SL1 region, there are no known single nucleotide variants with >1% in frequency and no known mutations among novel variants, this mechanism may represent a unique therapeutic target for immune-evasive, increasingly infectious strains that continue to emerge with the ongoing pandemic (Navarro Gonzalez et al., 2021). The recent appearance of variants as well as the history of spontaneous resistance of viruses to conventional antiviral drugs – most notoriously to nucleoside inhibitors – suggests the need for new classes of antivirals (Harvey et al., 2021; Seley-Radtke and Yates, 2018). The high conservation of both Nsp1 and SL1 and their requirement for viral replication suggest that SARS-CoV-2 mutants refractory to anti-SL1 ASO binding would be at a considerable selective disadvantage. This trade off can be exploited by anti-SL1 therapy, which could thus represent a potent antiviral strategy whose evasion is evolutionarily constrained, requiring co-mutation of SL1 and Nsp1. More generally, our proof-of-principle in developing therapeutics to unleash a pathogen’s own virulence mechanism upon itself may represent an important strategy to avoid resistance in SARS-CoV-2 and could be expanded to other host-pathogen systems.

## MATERIALS AND METHODS

### Plasmids and transfection

SARS-CoV-2 full-length Nsp1 (1-180 aa), Nsp1-NT (1-127 aa) and Nsp1-CT (128-180 aa) were amplified from pDONR207 SARS-CoV-2 NSP1 (Addgene) by PCR and then cloned into pDB-His-MBP or BacMam pCMV-Dest plasmid. Full-length 265 nt 5′ UTR of SARS-CoV-2 was subcloned to replace the 5’ UTR of human CMV in the pLV-mScarlet vector using a Hifi one-step kit (Gibson Assembly, NEB). Full-length, SL1 alone or ΔSL1 5′ UTR of SARS-CoV-2 were cloned into pLV-mScarlet vector or pGL3 basic vector. All constructs were verified by sequencing. Cells were transiently transfected with indicated plasmids using FuGENE Transfection Reagent (Promega) or Lipofectamine 2000 (Thermo Fischer Scientific) according to the manufacturer’s instructions. ASO transfection dosages correspond to the concentration at the time of lipid complex formation.

### Cell culture

HEK293T (ATCC CRL-3216, female), HeLa cells (ATCC CCL2, female), and, and Vero E6 (ATCC CRL1586) were purchased from the American Type Culture Collection, and Expi293 cells (female) were from ThermoFisher. Cells were cultured in Dulbecco’s Modified Eagle medium (Gibco) or Expi293 Expression Medium (Gibco) supplemented with 10% fetal bovine serum and 1% penicillin/streptomycin (Gibco) at 37 °C with 5% CO_2_.

### Luciferase assay

Luciferase assays were performed as previously described (Yang et al., 2020). Briefly, HEK293T cells were transfected with reporter plasmid and various Nsp1 constructs using FuGENE Transfection Reagent (Promega). Luciferase assays were performed using the Dual Luciferase Assay System (Promega); β-Galactosidase activity was used as an internal control. Luciferase activity was measured using a CytoFluorplate 4000 Luminescence Microplate Reader (ABI).

### Immunofluorescence and quantitative microscopy

To visualize the expression of mScarlet and MBP-tagged SARS-CoV-2 Nsp1 (WT, CT, NT, Nsp1-linker1 and Nsp1-linker2), transfected HeLa or 293T cells were fixed with 4% formaldehyde for 10 minutes at room temperature, permeabilized with PBS + 0.25% Triton X-100, and blocked with 3% BSA. Cells were stained with mouse anti-MBP (NEB, 1:500), followed by washing and subsequent incubation with goat anti-mouse IgG, Alexa Fluor 488 (1:1000, Invitrogen) as well as Hoechst 33342 counterstain (Immunochemistry). Cells were transfected in glass bottom 96 well plates (3-5 replicate wells per treatment) followed by imaging using a Zeiss Axioskop-2 microscope or an ImageXpress Micro Confocal (Molecular Devices) using a 20X objective (S Plan Fluor, NA=0.45). Multiple sites in each well were imaged to ensure large sample size (2-9, depending on instrument availability; each site contained ∼100-200 cells). Image data were analysed and quantified using Image J (NIH) or MetaXpress software (Molecular Devices). Representative images from Figures 1-4 were collected on an Olympus Fluoview FV1000 point scanning confocal microscope using a 60X UPlan S Apo water immersion objective (NA=1.2).

### Quantification and statistical analysis

Transfection experiments for microscopy were performed using at least 3 replicates in separate wells. The luciferase assay in Figure 3F was done on lysate samples pooled from 2 wells of replicates per treatment, and 3 separate measurements were taken for each lysate sample. Transfection-infection experiments in Figure 5E which were done in duplicate in 24 well plates. All graphs were plotted and analysed with GraphPad Prism 5 software. p > 0.05 was considered statistically not significant, and the following denotations were used: ***p < 0.001, **p < 0.01 and *p < 0.05. Sample size for each experiment corresponds to least 3 replicates except for Figure 5D which used two replicates.

### In vitro infection with SARS-CoV-2

Vero E6 cells in 24 well plates were seeded at ∼80% confluency and were co-transfected with 250ng Nsp1 and 100 nM of various ASOs and incubated overnight. The next day, the media was discarded, and cells were infected with 0.1 or 0.5 MOI in low serum media and allowed to incubate 72 hours. Cells were washed in PBS and fixed with 4% formaldehyde followed by permeabilization with 0.1% Triton-X 100 and staining with antibodies to N and 488-conjugated secondary antibodies.

## ACKNOWLEDGMENTS

P.F. was supported by a Cancer Research Institute Irvington Postdoctoral Fellowship. L.W. was supported by funding from an NIH T32 grant (5T32AI007512-34). S.M.V was supported by a postdoctoral fellowship from the American Cancer Society (133083-PF-19-034-01-LIB). T.-M.F. was supported by funding from an NIH T32 grant (5T32HL066987-18 to L.E.S.) and by start-up funds from the Ohio State University Comprehensive Cancer Center. Microscopy experiments using the ImageXpress high content imaging system were performed at the Harvard Medical School Longwood ICCB Drug Screening Facility. We would also like to thank Jiazhi Li and the Richard Gregory lab for technical assistance, Alexander B Tong, and Sichen (Susan) Shao for discussions.

The authors declare no competing interests.

## Author contributions

L.W., M.S., and H.W. conceived the project. L.W., P.F., and M.S designed the constructs and carried out cloning. S.M.V. and M.S. performed cell-based assays. S.M.V., M.S., and Y.Z. performed luciferase assays under J.L.’s supervision. V.L. and S.M.V. performed SARS-CoV-2 transfection-infection assay under L.G.’s supervision. L.W. and S.M.V designed the ASOs. P.F. and L.W performed EMSA. S.M.V., L.W., P.F., M.S., T.-M.F., and H.W. wrote the manuscript with input from all the authors.

## SUPPLEMENTAL MATERIAL

**Figure S1.**
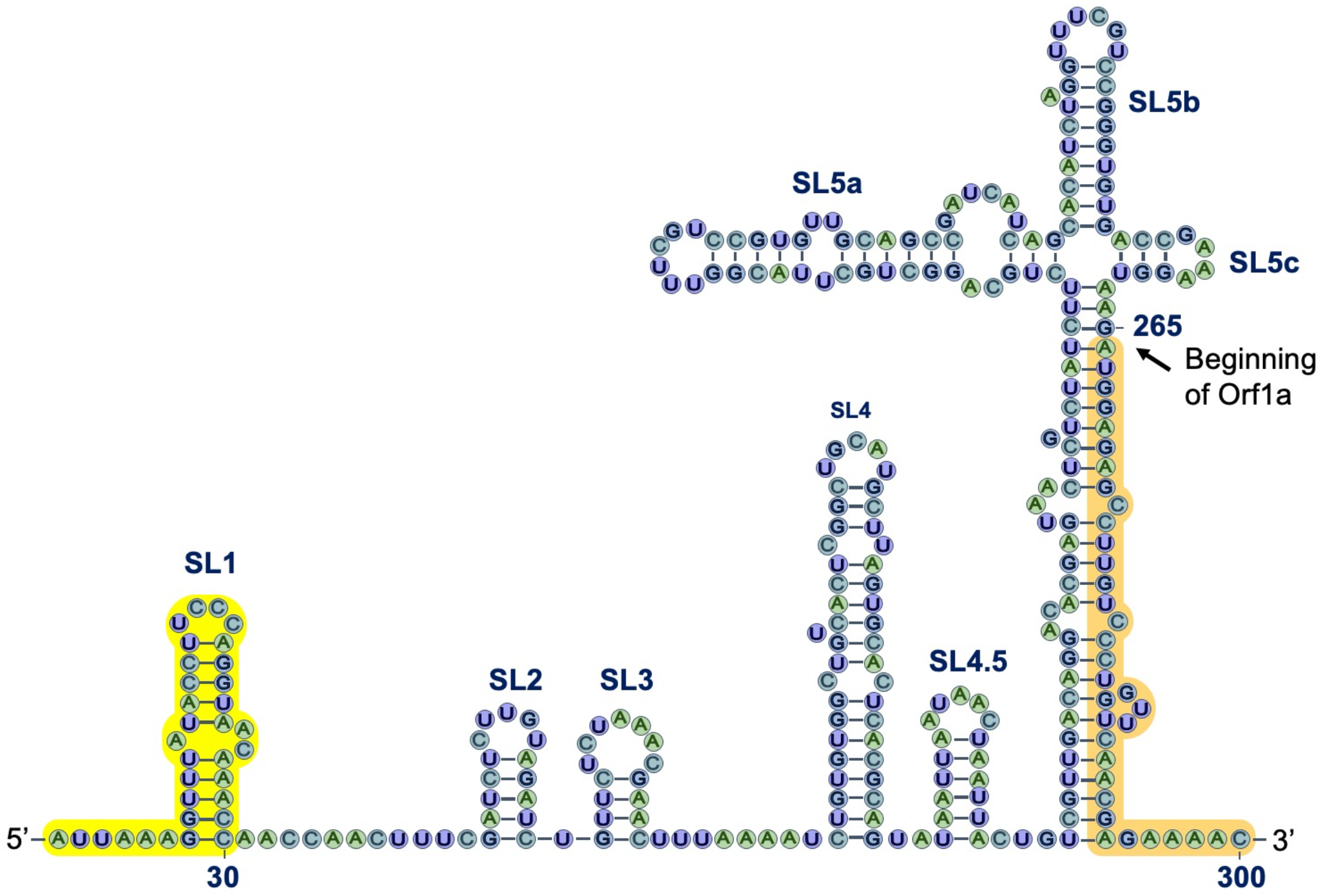
Schematic representation of SARS-CoV-2 5’ UTR secondary structure prediction. SL1 highlighted in yellow, and ORF1a highlighted in orange.

**Figure S2.**
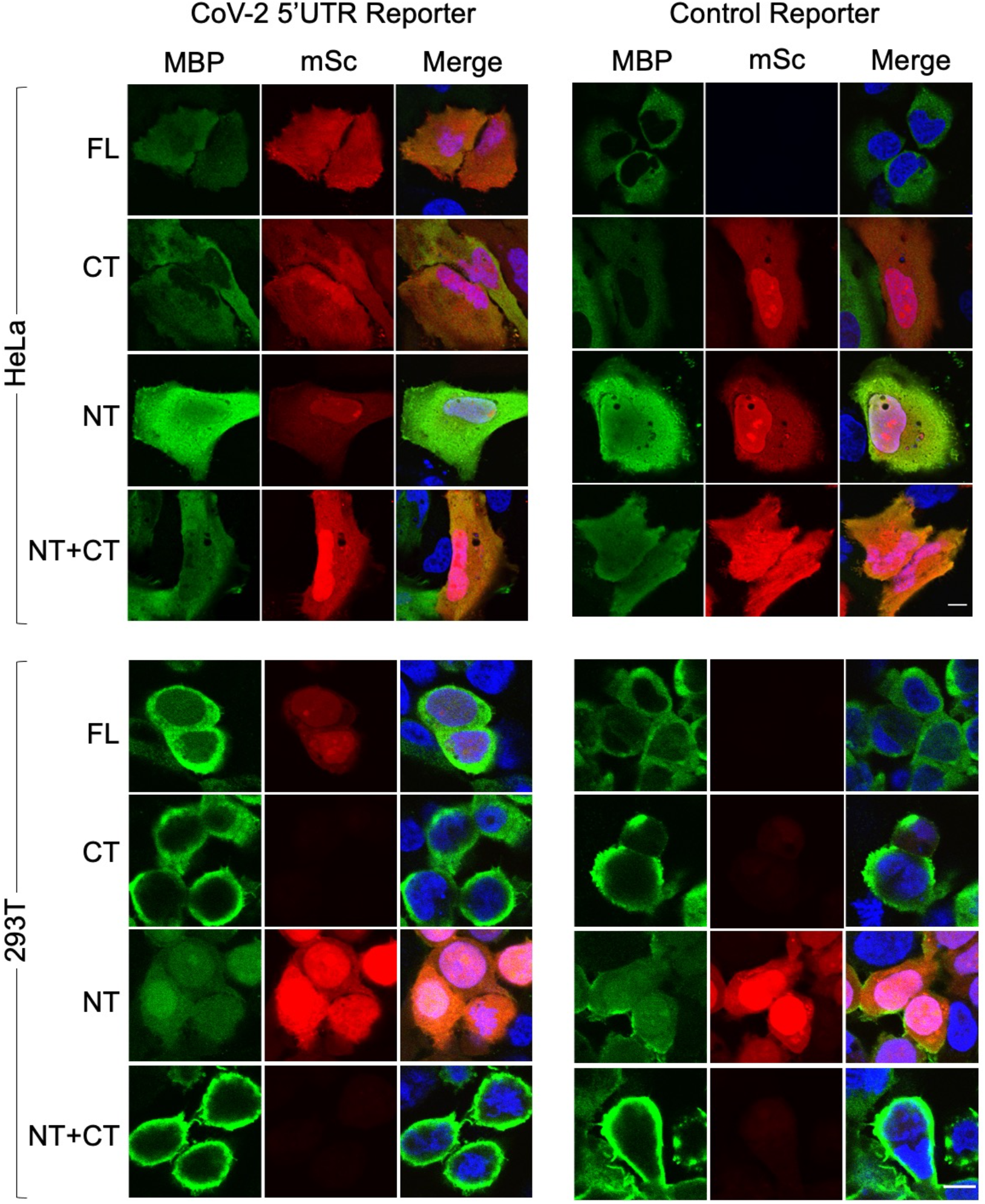
Immunofluorescence of MBP-Nsp1 and in situ fluorescence of reporter activity in HeLa and 293T cells. Cells were stained and visualized for various Nsp1 fragments tagged with MBP. Reporter activity was visualized with in situ mScarlet intensity. In the NT+CT condition, both NT and CT were tagged with MBP. Cells were counterstained with Hoescht 33342 to visualize nuclei.

**Figure S3.**
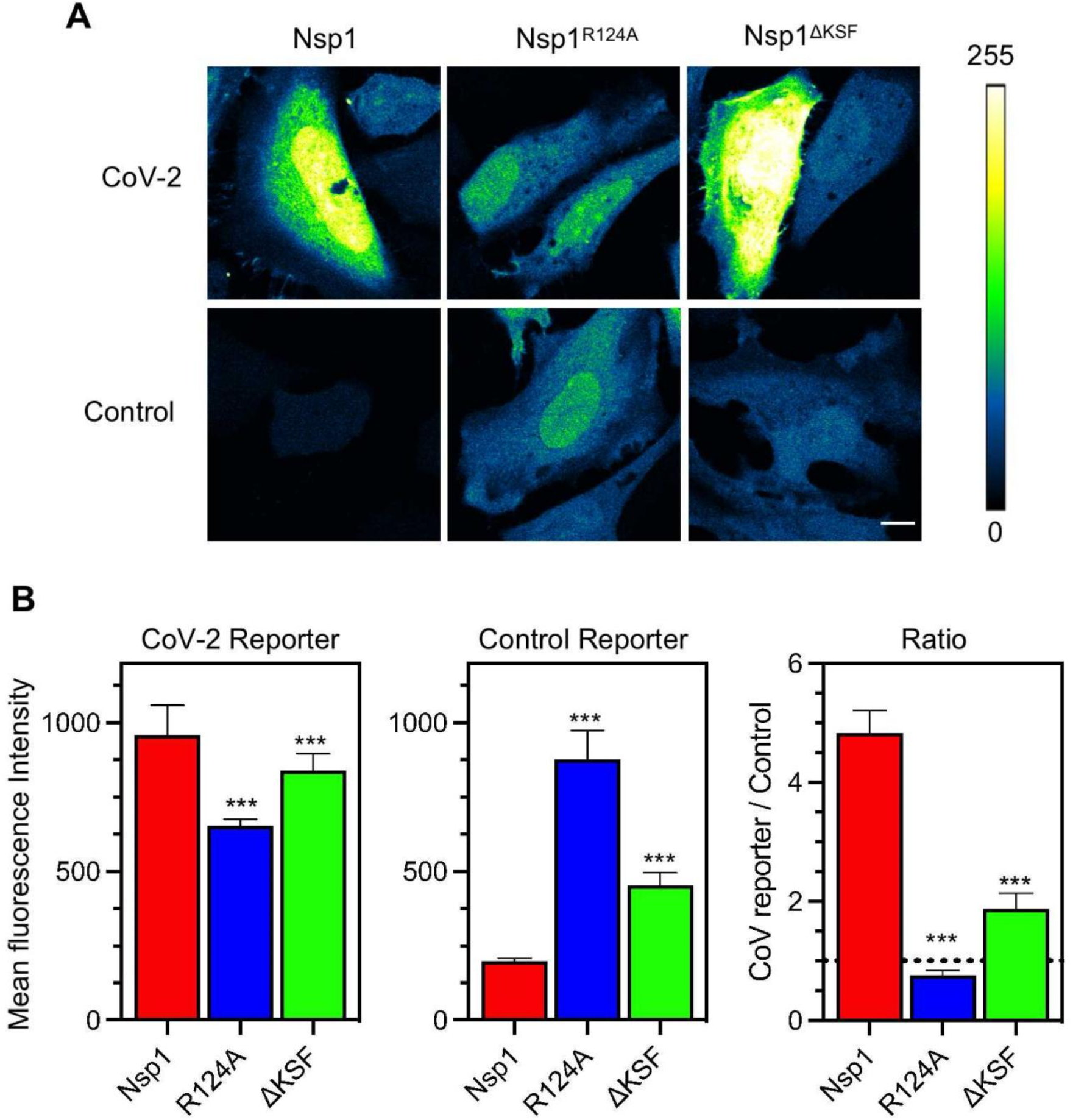
Nsp1^R124A^ and Nsp1^ΔKSF^ are defective in viral translation selectivity. (A) Images of reporter intensity in HeLa cells transfected with either CoV-2 or control reporter along with various mutants of Nsp1. (B) Mean fluorescence intensity of CoV-2 reporter or control reporter when co-transfected with Nsp1, Nsp1^R124A^, or Nsp1^ΔKSF^ (left and middle panels) and the ratio between the two reporters (right panel). Error bars represent standard deviation.

**Figure S4.**
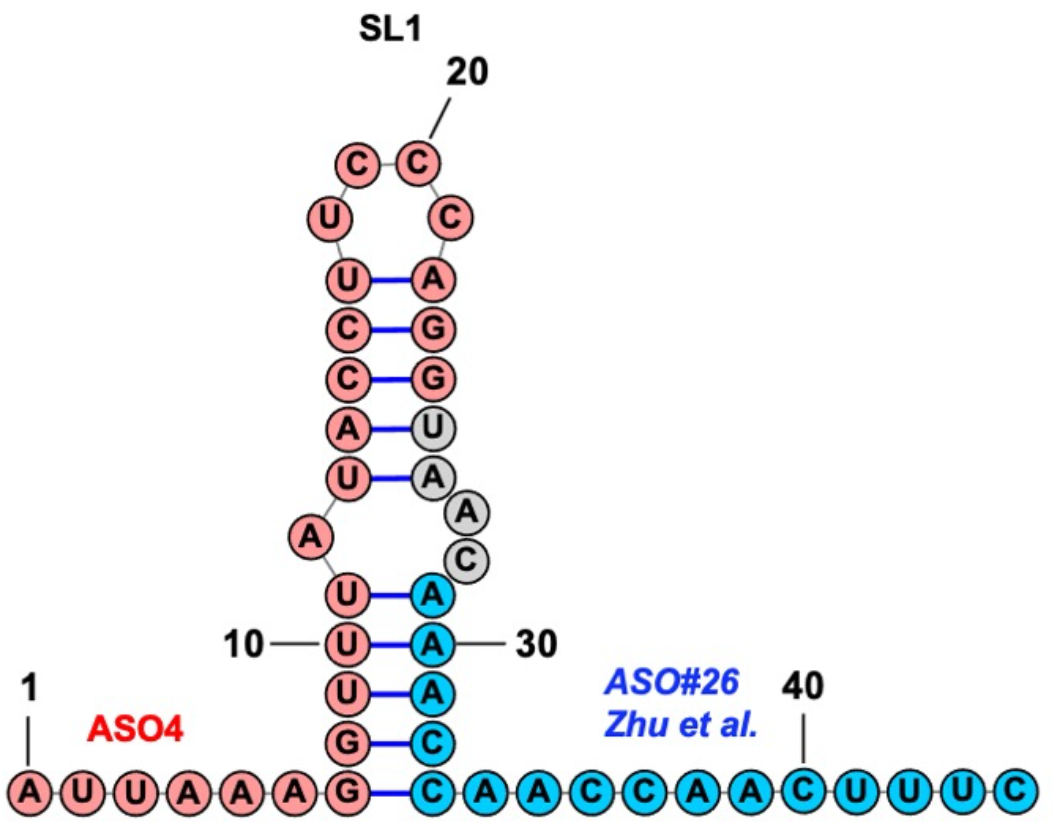
ASO4 (pink) and the ASO in (Zhu et al., 2021) (blue) target different regions of the 5’ UTR.

